# Early myelination involves the dynamic and repetitive ensheathment of axons which resolves through a low and consistent stabilization rate

**DOI:** 10.1101/2022.07.17.500324

**Authors:** Adam R. Almeida, Wendy B. Macklin

**Affiliations:** Department of Cell and Developmental Biology, University of Colorado School of Medicine, Aurora, CO 80045

## Abstract

Oligodendrocytes in the central nervous system exhibit significant variability in the number of myelin sheaths that are supported by each cell, ranging from just a few to about 50 (1–4). This begs the question, why does sheath number vary so widely across individual oligodendrocytes? Myelin production during development is dynamic and involves both sheath formation and loss (5–9). However, how these parameters are balanced to determine overall sheath number has not been thoroughly investigated. To explore this question, we combined extensive time-lapse and longitudinal imaging of oligodendrocytes in the developing zebrafish spinal cord to quantify sheath initiation and loss. Oligodendrocytes repetitively ensheathed the same axons multiple times before stable sheaths were formed. Interestingly, while there was a variable number of ensheathments produced by each oligodendrocyte, ~80-90% of these structures were always lost, which was an unexpectedly high, but consistent rate of sheath destabilization. The dynamics of this process indicated rapid membrane turn-over as ensheathments were formed and lost repetitively on each axon. To test whether membrane recycling impacts this sheath stabilization rate the endocytic pathway was disrupted by expressing a dominant-negative mutant form of Rab5. Oligodendrocytes over-expressing this mutant lost an even higher percentage of sheaths. Overall, oligodendrocytes initiate a highly variable number of axonal ensheathments, but these cells all exhibit a similar sheath stabilization rate, which is dependent on the endocytic recycling pathway.

## Introduction

Myelin sheaths are specialized cellular structures produced by oligodendrocytes in the central nervous system (CNS) that wrap around axons to accelerate the velocity of action potential conduction and to provide trophic support to neurons (reviewed in (10, 11)). The number of myelin sheaths that are supported by each oligodendrocyte is remarkably variable (reviewed in (12, 13)). Experiments using genetically encoded fluorescent reporters and immunostaining in both murine and zebrafish models establish that oligodendrocytes have anywhere from a few sheaths per cell up to about 50 (1–5, 14). However, the importance of this heterogeneity in sheath number and how it is created is still unclear.

Time-lapse imaging in zebrafish has shown that the dynamics of developmental myelination include both sheath production and sheath loss (5–9). As past studies have not thoroughly compared sheath formation and loss across the same individual cells, the question remains as to the relative contribution of these two mechanisms to establishing oligodendrocyte sheath number. Understanding this balance is important because there are dramatically different cellular and molecular mechanisms that regulate sheath formation and loss. Thus, both processes could potentially be manipulated to improve myelin regeneration.

To address this gap in our knowledge we combined extensive time-lapse and longitudinal imaging in the developing zebrafish spinal cord to quantify both sheath initiation and loss from the same cells. Unexpectedly, oligodendrocytes repetitively formed and retracted ensheathments from the same axon multiple times before any stable sheaths were formed. These dynamics resulted in all oligodendrocytes consistently stabilizing only ~10-20% of the total number of immature ensheathments produced by each cell. Altering the endocytic recycling pathway in oligodendrocytes further decreased overall sheath stability. These studies suggest that the proportion of ensheathments formed and lost is similar across all oligodendrocytes and that the endocytic recycling pathway regulates this process.

## Results

### Regional differences in oligodendrocyte sheath number during developmental myelination

The number of sheaths produced by a single oligodendrocyte in the zebrafish spinal cord at 4 days post fertilization (dpf) is widely variable (1). However, it is unclear how this variability arises during development. Oligodendrocytes in the spinal cord myelinate either dorsal or ventral axon tracts and occupy these regions with differing cell densities (Figure 1A and B). These domains have different functions, with ascending dorsal axons primarily transmitting sensory information and descending ventral axons transmitting motor information (15). We investigated whether some of the variability in oligodendrocyte sheath number comes from quantifying cells from both regions together. Zebrafish embryos were injected at the single-cell stage with a *sox10:eGFP-CAAX* plasmid to sparsely label the membrane of individual oligodendrocytes. At 4 dpf, sheath number ranged from 4-21 per cell (Figure 1C and D), which is in line with previous work (1). Importantly, oligodendrocytes in the dorsal region of the spinal cord had more sheaths on average than oligodendrocytes in the ventral region (Figure 1E). No statistically significant differences in average sheath length or the total sheath length were found when comparing these groups (Figure 1F and 1G). To investigate how the regional difference in sheath number is generated, we compared sheath initiation and loss by oligodendrocytes in the dorsal and ventral regions of the spinal cord.

**Fig. 1.**
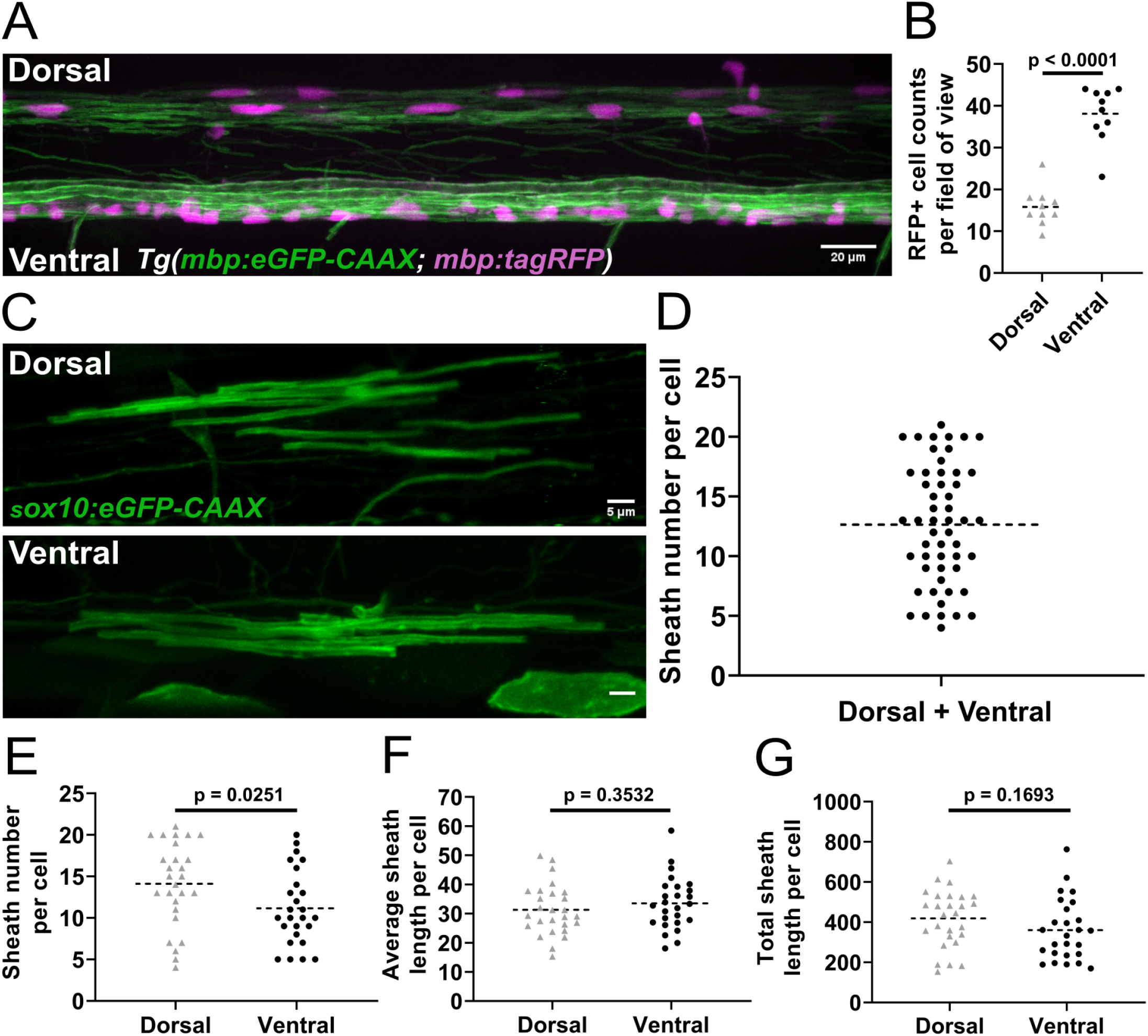
Sheath number was highly variable, but oligodendrocytes in the dorsal region of the spinal cord made more sheaths per cell than oligodendrocytes in the ventral region. **A**) Lateral image of the spinal cord of a living *Tg(mbp:eGFP-CAAX; mbp:TagRFP*) larva. (Scale bar = 20 μm). **B**) RFP+ oligodendrocyte cell counts in the dorsal and ventral regions of the spinal cord. (n = 10 larvae) **C**) Representative lateral images of dorsal and ventral oligodendrocytes in the spinal cord of living larvae at 4dpf labeled by *sox10:eGFP-CAAX* (Scale bar = 5 μm). **D**) Sheath number per cell, all data combined (n=53 cells/53 larvae). **E-G**) Sheath number per cell (E), average sheath length per cell (F), total sheath length per cell (G), comparing the dorsal and ventral regions (dorsal n = 27 cells/27 larvae, ventral n = 26 cells/26 larvae). The dashed lines in each plot represent average values with all data points shown. Significance determined by Mann–Whitney tests.

### The dynamics of early-stage sheath initiation and loss on individual axons

To establish parameters for quantifying the dynamics of sheath initiation and loss, we sparsely labeled neurons by injecting the pan-neuronal *neuroD:tagRFP-CAAX* axon reporter into *Tg(nkx2.2:eGFP-CAAX*), a transgenic myelin reporter line. The earliest stages of eGFP-CAAX-labeled sheath formation on tagRFP-CAAX-labeled axons were captured with extensive time-lapse imaging from 2.5-3dpf (Figure 2A, see methods for greater details). We could consistently identify immature ensheathments that were ~2μm or longer, surrounding axons with a cylindrical shape (Figure 2B and C). Of 68 ensheathments visualized, ~79% were lost by the end of the imaging period, which was an unexpectedly high rate of sheath destabilization. However, interestingly, this was because each oligodendrocyte repetitively ensheathed the same axonal domain an average of ~3 times before stable sheaths were formed (Figure 2D [T0’, T15’, T45’, T60’] and E). Repetitive ensheathments were multiple rounds of sheath initiation/loss on the same domain of an axon. Each axon domain was defined as the region between the two most lateral ensheathment attempts made by the same oligodendrocyte during a series of repetitive ensheathments. In cases with only a single ensheathment attempt, the axon domain was the region directly underneath the sheath. Of the 25 axonal domains analyzed, 14 of them ended with a stabilized sheath (Figure 2F) and 11 were destabilized (Figure 2G). Independent of outcome, the number of ensheathment attempts on each axonal domain ranged from 1-6. While there is obvious variability in the number of ensheathment attempts made on each axon, it is intriguing that 72% of the analyzed axonal domains were repetitively ensheathed more than once. This repetitive ensheathment phenomenon appears both common and fundamental to the process of building myelin sheaths.

**Fig 2.**
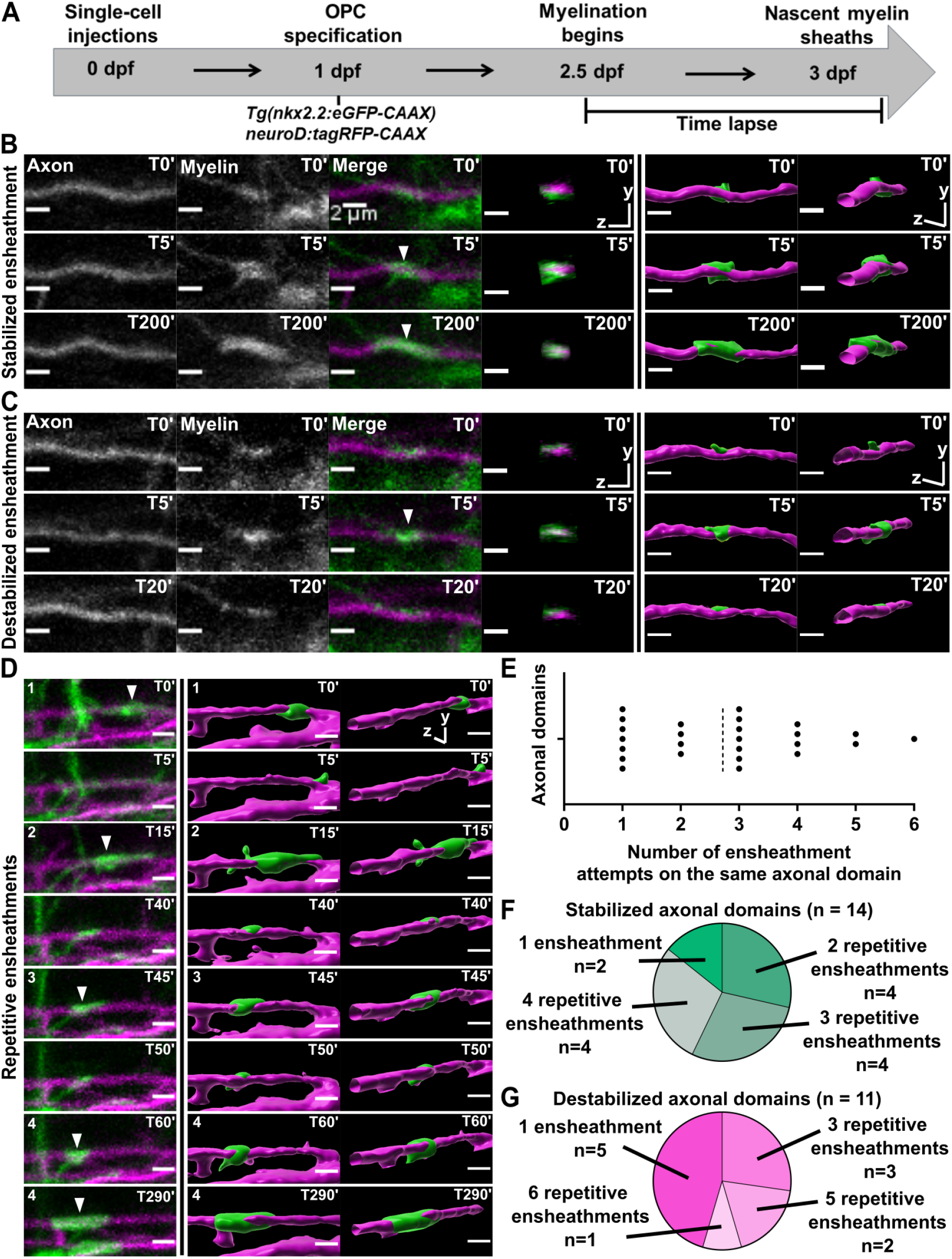
Oligodendrocytes repetitively ensheathed the same axonal domains before stable sheaths were formed. **A**) Axonal ensheathment dynamics experimental paradigm. **B-C**) Representative lateral images of immature ensheathments forming in the spinal cord of living *Tg(nkx2.2:eGFP-CAAX*) (myelin in green) larvae time-lapsed for 18 hours with a 5-minute imaging interval starting at 2.5dpf. Axons were labeled with *neuroD:tagRFP-CAAX* (magenta). YZ images are cross sections of the axon (volume projections from Imaris). 3D reconstructions were generated in Imaris. **B**) Time-lapse imaging of a stabilized ensheathment (Scale bar = 2 μm). **C**) Time-lapse imaging of a destabilized ensheathment (Scale bar = 2 μm). **D**) Representative lateral images of an axonal domain that was repetitively ensheathed 4 times before a sheath was stabilized. 3D reconstructions were generated in Imaris (Scale bar = 2 μm). **E**) Plot showing the number of times each axonal domain was ensheathed, e.g., 7 axonal domains had only 1 ensheathment attempt. Dotted line represents the average number of ensheathment attempts. (n = 25 axonal domains, 18 axons, 12 larvae, 68 ensheathments imaged total) **F**) Pie chart of the number of axonal domains with a final stable ensheathment and the number of ensheathment attempts that were made on each of those domains (n = 14 axonal domains). **G**) Pie chart of the number of destabilized axonal domains and the number of ensheathment attempts that were made on each of those domains (n =11 axonal domains).

### Oligodendrocyte nascent sheath accumulation was highly dynamic

To capture the overall dynamics of the ensheathment process in the context of individual oligodendrocytes, we modified the ensheathment dynamics imaging paradigm from Figure 2 (Figure 3A). Embryos were co-injected with our *sox10:eGFP-CAAX* oligodendrocyte lineage cell membrane reporter and *myrf:tagRFP*, a cytosolic reporter expressed in myelin-fated oligodendrocytes. This allowed us to identify sparse, double-labeled eGFP^+^RFP^+^ cells in both the dorsal and ventral tracts of the spinal cord at 2.5 dpf. We captured the cellular dynamics of each cell for 15 hours with a 5-minute imaging interval. At 3 dpf, larvae were placed back into embryo medium. At 4dpf, the same larvae were re-mounted, and a single static image was taken for each of the same cells. From this, all sheath initiations and losses for each cell could be tracked, and quality control experiments determined that these imaging conditions did not change overall sheath number or average sheath length (Figure 3.1 supp, see methods for further details).

**Fig 3.**
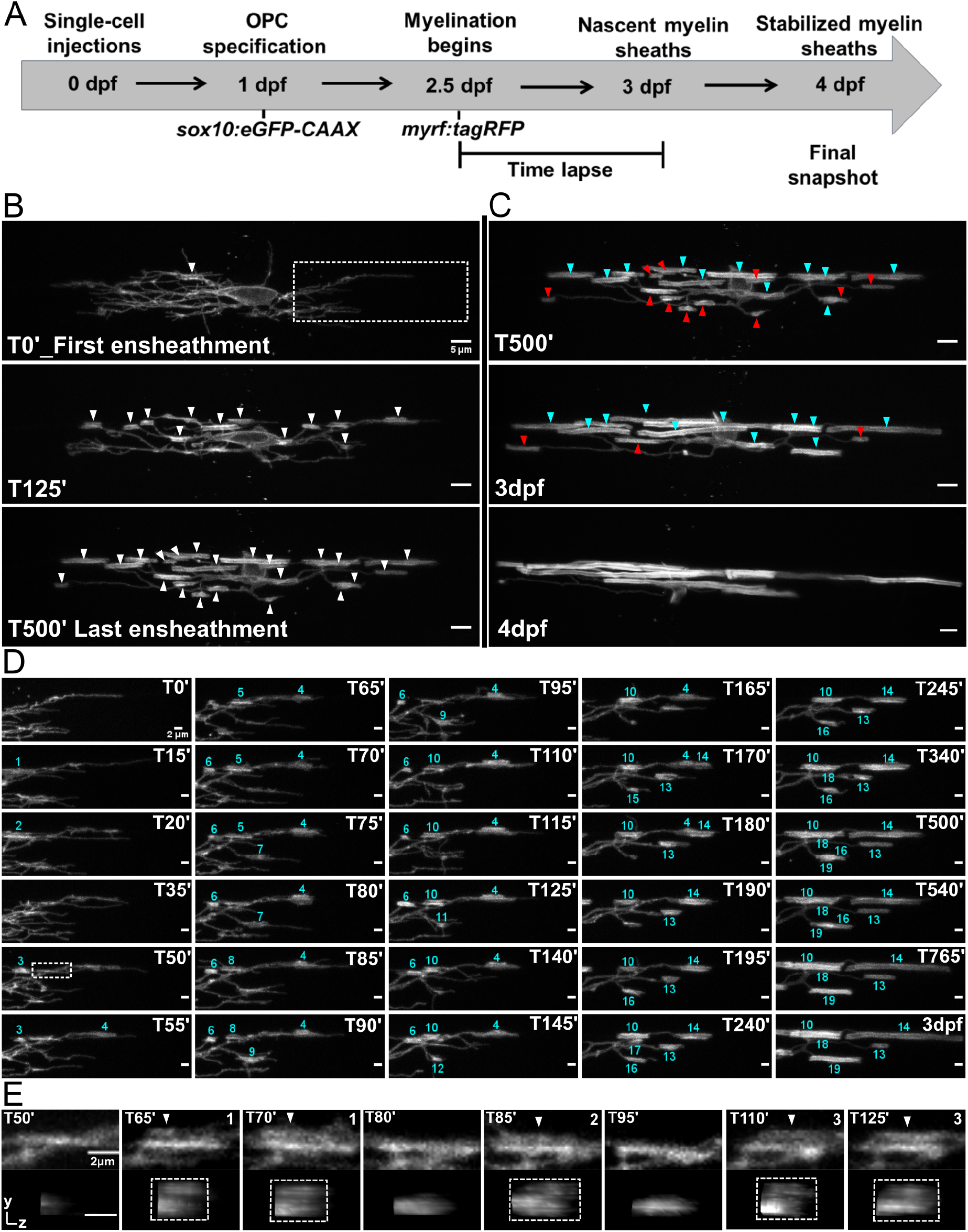
Nascent sheath accumulation involved frequent and repetitive sheath initiation and loss. **A**) Oligodendrocyte ensheathment dynamics imaging paradigm. **(B-E**) Lateral images of oligodendrocyte ensheathment dynamics in the spinal cord of living larvae labeled with *sox10:eGFP-CAAX* and time-lapsed for 15 hours with a 5-minute imaging interval from 2.5-3dpf. **B**) Images show the progression of immature sheath accumulation from the first ensheathment attempt in the first panel (set at T0’) to the final ensheathment at T500’ in the bottom panel. Outlined region in first panel is analyzed further in Figure D and E. White arrows identify all immature ensheathments in each frame. (Scale bar = 5 μm). **C**) Images showing sheath loss during the stabilization phase. T500’ image is presented again in the top panel and is relabeled to identify future stabilized (cyan) or destabilized (red) ensheathments. This oligodendrocyte has 22 immature sheaths at T500’, 11 of which disappeared, and it has 11 stabilized sheaths at 4dpf in the final panel (Figure 3.2 supp). (Scale bar = 5 μm). **D**) Images of the outlined region in the first panel of Figure B visualizing frequent sheath initiation and loss. Each ensheathment is represented with the numbers 1-19 to signify the order that each one appears. The numbers disappear when an ensheathment is lost. Only 5 ensheathment attempts were stabilized out of a total of 19. (Scale bar = 2 μm) **E**) Further enlarged images of the outlined region in Figure D (T50’) demonstrate the same repetitive ensheathment phenomena as in Figure 2. White arrows point out 3 ensheathment attempts before the final one was stabilized. The YZ images are cross sections of each ensheathment (volume projections in Imaris). A box is drawn around each cylindrical ensheathment in the YZ images. (Scale bar = 2 μm).

To begin answering the question of how sheath initiation and loss is balanced for individual oligodendrocytes, these events were manually tracked and quantified in each frame of our time-lapse data set. As in published studies (5), we found that oligodendrocytes have an initial phase of immature sheath accumulation (Figure 3B), followed by a phase of sheath stabilization and loss (Figure 3C). Interestingly however, the accumulation phase was very dynamic in our studies, with a surprisingly high rate of sheath destabilization. For example, only 5 out of 19 ensheathment attempts were stabilized for the outlined region of the cell in Figure 3B (Figure 3D). Even without directly imaging axons, we readily observed instances of presumed repetitive ensheathment (Figure 3E).

To better understand the impact of these repetitive ensheathment dynamics, we needed to average the levels of sheath initiation and loss across all cells in our data set and establish a graphic method for presenting the results. However, the number of hours over which each cell accumulates sheaths ranged from ~4-8 hours (Figure 4A), making it difficult to combine data from multiple cells for quantitative comparison. Nevertheless, we identified a definable transition point for each cell from accumulating new immature sheaths to stabilizing them. Thus, once an oligodendrocyte accumulated a peak (or max) number of immature sheaths, every cell slowed down and stopped forming new sheaths. Given the common dynamic of accumulation and plateau, it was therefore possible to normalize the timing of sheath accumulation in each video by defining the frame at which each cell had accumulated its peak number of sheaths as time zero for that cell (Figure 4B). Using this normalization for 19 individual dorsal cells and 18 individual ventral cells, we found that the dorsal cells accumulated a higher peak number of ensheathments compared to the ventral cells (Figure 4B). Additionally, ~95% of all ensheathment attempts occurred before reaching this peak for both groups (Figure 4C and Figure 4D vertical dashed line). This was an average of ~100 ensheathment attempts for dorsal cells and ~74 for ventral cells. Interestingly, ~75% of these ensheathments were also lost before reaching peak for both groups (Figure 4E and F). Collectively, this data shows that oligodendrocytes initiate a remarkable number of immature ensheathments, with only a small percentage maintained by the end of the accumulation phase.

**Fig 4.**
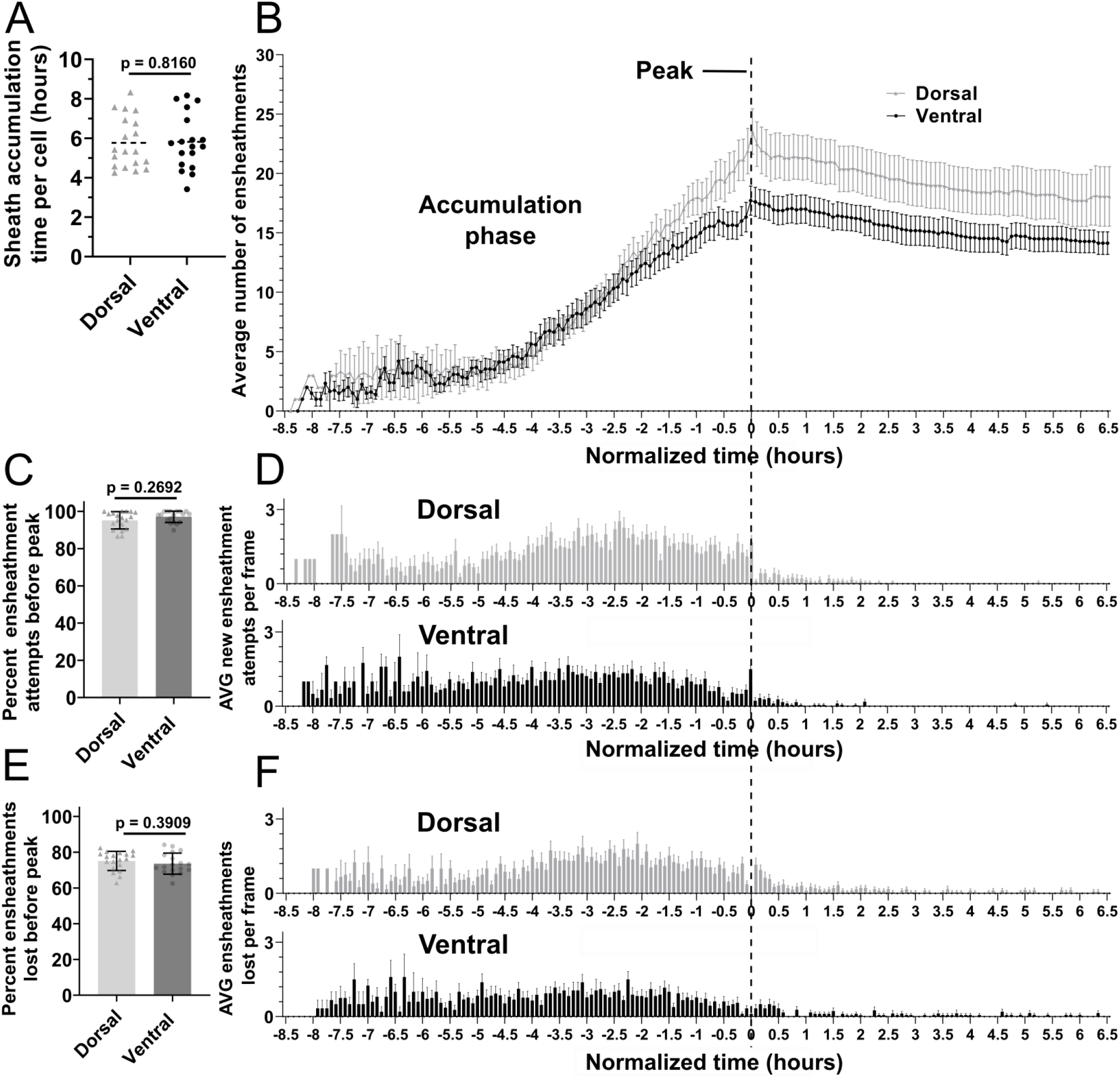
Oligodendrocyte nascent sheath accumulation involved a consistent and high rate of sheath initiation and loss. **A**) Time required for peak (or max) sheath accumulation per cell (hours). **B-F**) Videos of n = 19 dorsal cells (19 larvae) and n = 18 ventral cells (18 larvae) were quantified for sheath initiation and loss in each 5 min frame. The time-lapse videos were then normalized for quantitative comparison by defining the video frame at which each cell accumulated its peak (or max) number of immature ensheathments as T0’. Each frame before or after that was -5 min or +5 min. The vertical dashed line aligns the T0’ time point of each graph in B, D, and F. **B**) The average number of immature ensheathments for dorsal and ventral cells that are present in each frame is plotted based on time relative to T0’; error bars represent SEM. **C**) Percent of the total number of ensheathment attempts that occurred prior to reaching peak. Error bars represent SEM. **D**) The average number of new ensheathment attempts in each frame is plotted with the same time normalization as in B. Error bars represent SEM. **E**) Similar to C, the percent of the total number of ensheathments that were lost prior to reaching peak. Error bars represent SEM. **F**) The average number of ensheathments that were lost in each frame is plotted with the same time normalization as in B. Error bars represent SEM. Individual data points are shown and significance was determined by Mann–Whitney tests in A, C, and E. The data in B, D, and F was cropped at +6.5 hours.

### All oligodendrocytes exhibit a low and consistent sheath stabilization rate

It is very striking that all oligodendrocytes exhibited a similar rate of sheath destabilization and loss during the accumulation phase (Figure 4), yet dorsal cells end up with more sheaths than ventral cells. Dorsal oligodendrocytes exhibited more ensheathment attempts, accumulated a higher peak number of immature ensheathments, and maintained more sheaths at the 4 dpf time point compared to ventral cells (Figure 5A-E). Dorsal oligodendrocytes also lost more sheaths during the stabilization phase compared to ventral cells (Figure 5F). Surprisingly, however, both populations stabilized a similar percentage of ensheathments throughout the accumulation, stabilization, and combined phases of our experimental paradigm (Figure 5G-I). Thus, the levels of sheath initiation for each cell are very heterogeneous (ranging from ~20-190), but all oligodendrocytes exhibit a similar overall rate of sheath stabilization (~10-20%). To be confident with our quantification of this stabilization rate, we needed to be sure that these cells do not produce any new sheaths from 3-4dpf during the sheath stabilization phase. We designed a modified ensheathment dynamics imaging paradigm to test this and found that oligodendrocytes made no new sheaths during this time interval (Supp figure 5.1, see methods for further details). Collectively, we conclude that dorsal cells end up with more sheaths at 4dpf compared to ventral cells due to increased sheath initiation.

**Fig. 5.**
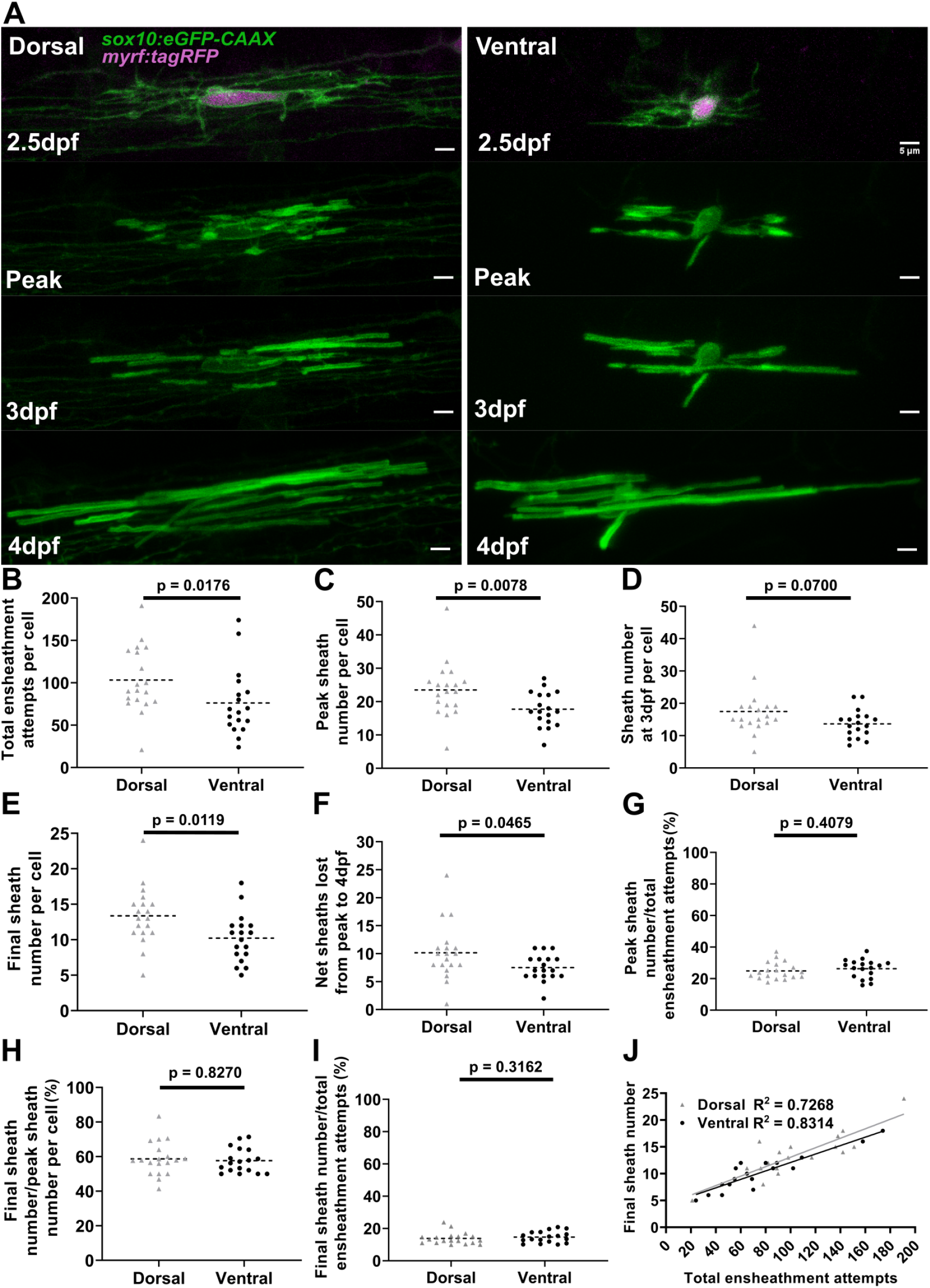
Dorsal cells initiated more ensheathments than ventral cells, but all oligodendrocytes stabilized a similar percentage of these ensheathments. **A**) Dorsal and ventral cell lateral images from the ensheathment dynamics imaging paradigm in the spinal cord of living larvae labeled with *sox10:eGFP-CAAX* (in green) and time-lapsed for 15 hours from 2.5-3dpf. The upper panels are a dorsal (left) and ventral (right) cell also labeled with *myrf:tagRFP* (magenta) at the beginning of the time-lapse experiment. The subsequent panels are the same cells at the peak of sheath accumulation, at 3dpf, and at 4dpf. (Scale bar = 5 μm). **B-J** compares dorsal and ventral cells. **B**) Total ensheathment attempts per cell. **C**)Peak sheath number per cell. **D**) Sheath number at 3dpf per cell. **E**) Final sheath number per cell at 4dpf (This is the same data and images as presented in supplemental **Figure 3.1**, time-lapse group). **F**) Net sheaths lost from the peak to 4dpf. **G**) Percent of sheaths stabilized during the accumulation phase (peak sheath number/total ensheathment attempts). **H**)Percent of sheaths stabilized during the stabilization phase (final sheath number/peak sheath number). **I**) Percent of total sheaths stabilized across both the accumulation and stabilization phases (final sheath number/total ensheathment attempts). **J**) Simple linear regression comparing the total number of ensheathment attempts to the final sheath number at 4dpf for each cell. The R^2^ values for each group are shown. Dorsal n=19 cells/19 larvae, ventral n=18 cells/18 larvae. The dashed lines in each plot represent average values with all data points shown. Significance was determined by Mann–Whitney tests.

We predict that this consistent sheath stabilization rate could be a fundamentally important parameter for understanding oligodendrocyte ensheathment behavior. Indeed, a simple linear regression comparing the total number of ensheathment attempts with the final number of sheaths for each cell established high correlation of these parameters (R^2^ value of ~0.73 for dorsal cells and ~0.83 for ventral cells, Figure 5J). Thus, for each stabilized ensheathment, there is a relatively predictable number of ensheathments that are destabilized and lost. Altogether, these results strongly suggest that both sheath initiation and loss are proportionately regulated across all oligodendrocytes.

### Components of the endocytic recycling pathway localized to immature sheaths and regulated myelin sheath number

The repetitive ensheathment of axons is orchestrated by a dynamic series of morphological transitions that are energetically expensive for oligodendrocytes. We hypothesized that the endocytic recycling of membrane and membrane proteins has a role in these transitions (Depicted in Figure 6A), and we therefore tested the impact of interfering with the endocytic recycling pathway on the ensheathment dynamics of oligodendrocytes.

**Fig. 6.**
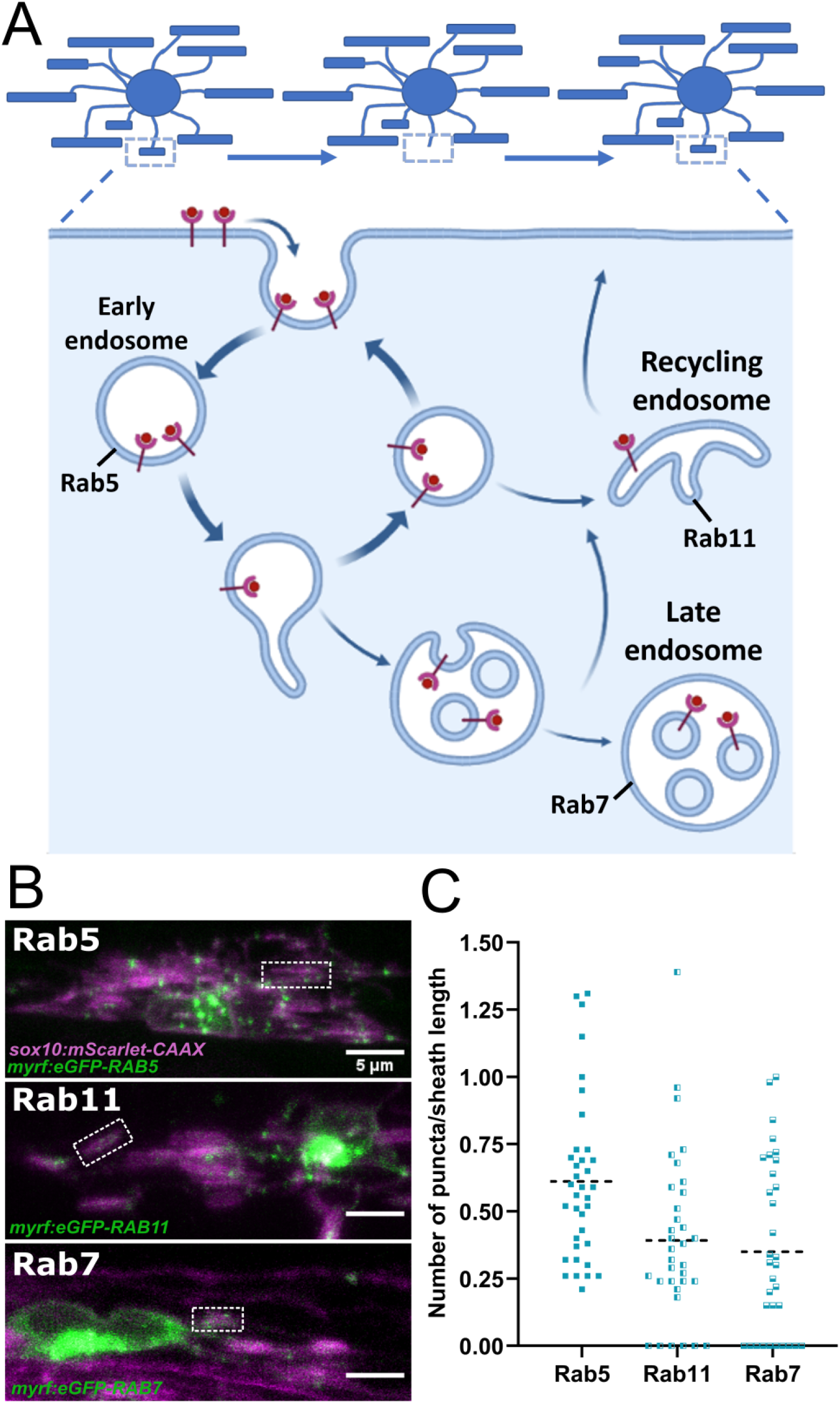
Rab5+, Rab7+, and Rab11+ endosomes localized to immature sheaths. **A**) The endocytic recycling pathway during sheath initiation and loss (Adapted from “Endocytic Pathway with Macropinocytosis and Phagocytosis”, by BioRender.com (2022). Retrieved from https://app.biorender.com/biorender-templates/t-5ea05f35722d6800ab456324-endocytic-pathway-with-macropinocytosis-and-phagocytosis). **B**) Lateral images of oligodendrocytes in the early stages of the ensheathment process in the spinal cord of living larvae at 2.5 dpf labeled with *sox10:mScarlet-CAAX* (magenta) and expressing either *myrf:eGFP-RAB5C, myrf:eGFP-RAB7A*, or *myrf:eGFP-RAB11A* (green). White boxes outline immature sheaths with Rab+ endosomal puncta for each fusion protein. (Scale bar = 5 μm). **C**) Quantification of Rab+ endosomal puncta in immature sheaths. Number of puncta in each sheath was normalized by the length of the sheath. Rab5c n = 36 sheaths (9 ventral cells/2 dorsal cells/11 larvae), Rab11a n = 34 sheaths (8 ventral cells/2 dorsal cells/10 larvae, Rab7a n = 33 sheaths (9 ventral cells/2 dorsal cells/11 larvae). The dashed lines represent average values for all data points.

The endocytic recycling pathway is controlled by several Rab-GTPase proteins that act as master regulators of membrane trafficking (reviewed in (16)). Rab5, Rab7, and Rab11 regulate the early endosome, late endosome/lysosome, and recycling endosome respectively (Figure 6A). The cargo of endocytic vesicles is sorted at the early endosome (Rab5) and is then trafficked back to the plasma membrane or to the recycling endosome (Rab11) for further sorting. Alternatively, molecular cargo will be degraded as the early endosome matures into a late endosome/lysosome (Rab7) (reviewed in (17, 18)).

We initially analyzed whether the Rab5, Rab7 or Rab11 proteins localized to immature myelin sheaths. Expression constructs were generated to specifically label Rabs in oligodendrocytes, using *myrf* regulatory DNA to drive eGFP-Rab fusion proteins: *myrf:eGFP-RAB5C, myrf:eGFP-RAB7A*, and *myrf:eGFP-RAB11A*. We transiently expressed each Rab fusion protein with *sox10:mScarlet-CAAX* to label oligodendrocyte membranes. Images of early myelinating oligodendrocytes expressing these Rab proteins were collected at 2.5 dpf (Figure 6B), and the number of Rab+ puncta present in each immature sheath was counted and normalized by the length of each sheath. All 3 types of endosomes were present within immature sheaths (Figure 6C). This data suggests that Rab5, Rab7, and Rab11 could all potentially play a role in sheath formation.

To investigate whether Rab5, Rab7, or Rab11 are important for regulating sheath number in oligodendrocytes, we performed a sheath analysis in cells expressing dominant-negative Rab mutants (19, 20). Rab proteins are GTPases that are active when bound to GTP and inactive when bound to GDP (reviewed in (21)). Thus, the *rab5C^S36N^, rab7A^T22N^*, and *rab11A^S25N^* dominant negative point mutations maintain the Rab protein in a GDP bound ‘off’ state. However, since these mutant proteins still bind GEF proteins that normally activate Rabs, they reduce the level of GEFs available to activate endogenous Rabs, thereby decreasing the overall activity of the endogenous Rab protein.

We injected *myrf:tagRFP-rab5C^S36N^, myrf:tagRFP-rab7A^T22N^*, and *myrf:tagRFP-rab11A^S25N^* fusion constructs into embryos alongside *sox10:eGFP-CAAX* to label oligodendrocyte membranes. We captured static images of ventral cells only at 4 dpf and quantified myelin sheath number and sheath length. We found that the *rab5C^S36N^* and *rab11A^S25N^* mutants both decreased the average sheath number per cell without changing average sheath length (Figure 7A-D). Thus, expression of these mutants apparently had no general negative impact on oligodendrocyte development, but they likely regulate a specific aspect of the ensheathment process. On the other hand, the *rab7A^T22N^* mutant had little impact on sheath number or sheath length (Figure 7A-D), which suggests that the phenotype we observe for the *rab5C^S36N^* and *rab11A^S25N^* mutants is specific to the recycling pathway and is not a generic effect of expressing a dominant-negative Rab mutant.

**Fig 7.**
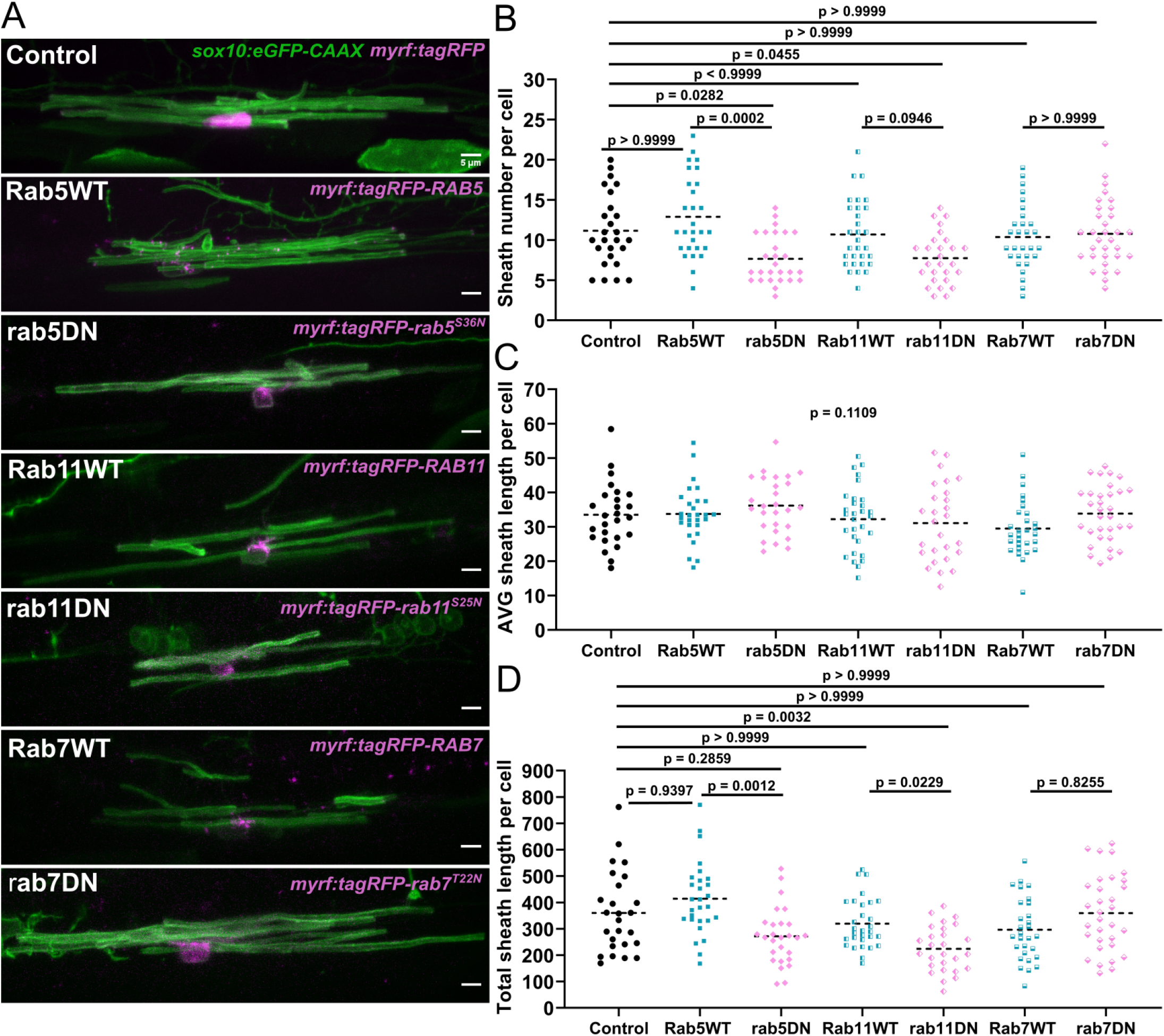
The *rab5C^S36N^* and *rab11A^S25N^* dominant-negative mutants reduced sheath number, but not average sheath length. **A**) Representative lateral images of ventral oligodendrocytes in the spinal cord of living larvae at 4dpf labeled by *sox10:eGFP-CAAX* (green) and one of the following: *myrf:tagRFP, myrf:tagRFP-RAB5C, myrf:tagRFP-rab5C^S36N^, myrf:tagRFP-RAB7A, myrf:tagRFP-rab7A^T22N^*, and *myrf:tagRFP-RAB11A, myrf:tagRFP-rab11A^S25N^* (all in magenta). The image and data for the control is the same as for the ventral group in Figure 1. (Scale bar = 5 μm). **B**)Sheath number per cell. **C**) Average sheath length per cell. **D**)Total sheath length per cell *(myrf:tagRFP* n=26 cells/26 larvae, *myrf:tagRFP-RAB5C* n=28 cells/28 larvae, *myrf:tagRFP-rab5C^S36N^* n=27 cells/27 larvae*, myrf:tagRFP-RAB7A* n=29 cells/29 larvae, *myrf:tagRFP-rab7A^T22N^*n=32 cells/32 larvae, and *myrf:tagRFP-RAB11A* n=30 cells/30 larvae, *myrf:tagRFP-rab11A^S25N^* n=27 cells/27 larvae). The dashed lines in each plot represent average values with all data points shown. Global significance was determined using a Kruskal-Wallis test for B-D. This global p-value is shown for C since it was not significant. Post hoc multiple comparison tests were not performed for this analysis. Post hoc Dunn’s multiple comparison tests were performed to compare groups in B and D. Before running the tests, we decided to compare everything with the control group and to also compare each wild-type and associated mutant with each other. The different Rab groups were not compared with each other.

### Rab5 regulates longitudinal sheath stability

We hypothesize that the repetitive ensheathment of axons is required to facilitate optimal sheath accumulation and is regulated by the endocytic recycling pathway. We therefore predict that disrupting endocytic recycling will reduce these dynamics and diminish overall sheath accumulation. Both Rab5 and Rab11 are involved in the recycling component of the endocytic pathway and their dominant negative mutants both decreased the overall sheath number per cell. Next we decided to specifically investigate Rab5 in depth. We analyzed cells expressing the *rab5c^S36N^* mutant in the ventral spinal cord tracts using the same ‘oligodendrocyte ensheathment dynamics paradigm’ described above. Oligodendrocytes expressing either *myrf:tagRFP-RAB5C* or *myrf:tagRFP-rab5C^S36N^* were imaged and compared to the ventral control data from Figures 3–5.

Unexpectedly, the over-expression of the *rab5c^S36N^* mutant did not change the dynamics of the accumulation phase. These cells formed and lost the same number of ensheathments during this phase and accumulated ensheathments to a similar peak number compared to the controls (Figure 8A-C, Figure 8.1 supp A-F). Instead, the *rab5C^S36N^* mutant specifically impacted the stabilization phase (which starts after cells reach peak sheath accumulation). We found a strong trend that the mutant increased the net number of sheaths that were lost from the peak sheath number out to 4dpf although it was not statistically significant (Figure 8C-F).

**Fig. 8.**
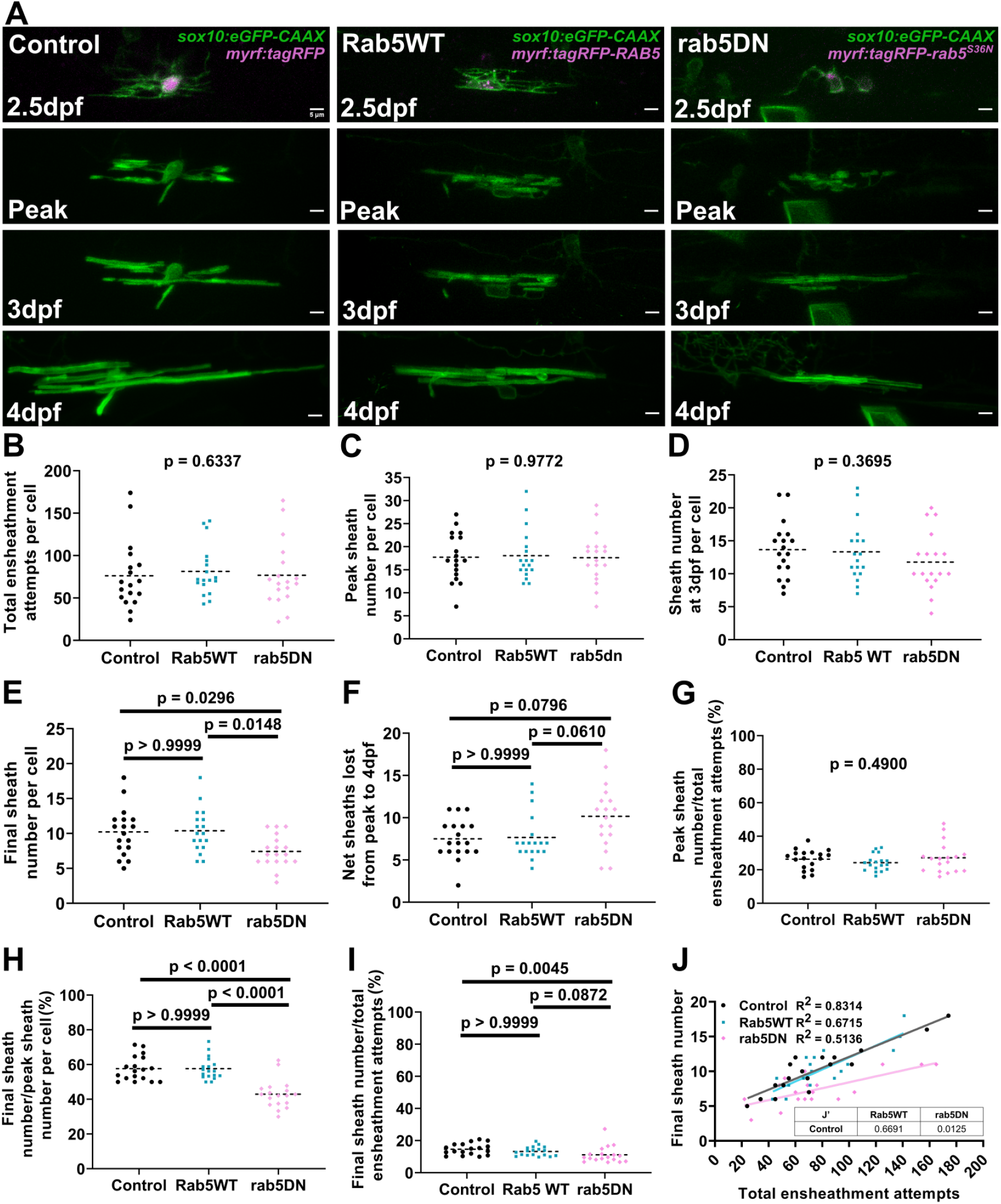
Over-expression of the *rab5C^S36N^* dominant negative mutant in oligodendrocytes reduced sheath stability in the stabilization phase. Lateral images from the ventral spinal cord of living larvae labeled with *sox10:eGFP-CAAX* (in green) and one of the following: *myrf:tagRFP, myrf:tagRFP-RAB5C, myrf:tagRFP-rab5C^S36N^* (all in magenta); and time-lapsed for 15 hours from 2.5-3dpf. The first panel is an image taken right before starting the time-lapse experiment. The subsequent panels are the same cells at the peak of sheath accumulation, at 3dpf, and at 4dpf. The images and data for the control are the same as for the ventral group in Figure 5. (Scale bar = 5 μm). **B**) Total ensheathment attempts per cell. **C**) Peak sheath number per cell. **D**) Sheath number at 3dpf per cell. **E**) Final sheath number per cell at 4dpf. **F**) Net sheaths lost from the peak to 4dpf. **G**) Percent of sheaths stabilized during the accumulation phase (peak sheath number/total ensheathment attempts). **H**) Percent of sheaths stabilized during the stabilization phase (final sheath number/peak sheath number). **I**) Percent of total sheaths stabilized across both the accumulation and stabilization phases (final sheath number/total ensheathment attempts). **J**) Simple linear regression comparing the total number of ensheathment attempts to the final sheath number at 4dpf for each cell. (control n=18 cells/18 larvae, wild-type *RAB5C* n=18 cells/18 larvae, *rab5C^S36N^* n=18 cells/17 larvae). The dashed lines in each plot represent average values with all data points shown. Significance was determined using global Kruskal-Wallis tests. These p-values are shown for B-D and G since they were not significant. Post hoc multiple comparisons tests were not performed for these analyses. Post hoc Dunn’s multiple comparisons tests were done for E, F, H, and I and the individual p-values are shown. **J’**)The slopes of the Rab5WT and rab5DN regression lines from J were compared to the control in Graphpad by (two-tailed) testing the null hypothesis that the slopes are identical (the lines are parallel). P-values are shown in the table.

Consistent with this trend, cells over-expressing the *rab5C^S36N^* mutant stabilized a lower percentage of sheaths in the stabilization phase specifically (Figure 8G-I). A simple linear regression comparing the total ensheathment attempts to the final sheath number showed that the slope of the mutant regression line was significantly lower than for the control group (Figure 8J). Collectively, we conclude that the dominant-negative *rab5C^S36N^* mutant expressed from the *myrf* driver negatively impacts the longitudinal sheath stabilization rate of oligodendrocytes but has little or no impact on initial sheath accumulation.

## Discussion

How is sheath initiation and loss balanced to regulate sheath number in oligodendrocytes? We addressed this fundamental question by combining time-lapse and longitudinal imaging to monitor the dynamics of this process from the same cells across time and demonstrate that it is the number of initial ensheathments produced by each cell that determines sheath number. However, this is because ~80-90% of all ensheathments produced by each cell are consistently lost. This high rate of destabilization is a result of repetitively ensheathing the same axons multiple times on average before stable sheaths are formed. We perturbed the cellular dynamics of the ensheathment process by disrupting the endocytic recycling pathway. Unexpectedly, we found that inhibiting the activity of Rab5, a GTPase protein that regulates trafficking at the early endosome, caused oligodendrocytes to lose more sheaths during the stabilization phase specifically. We therefore conclude that Rab5 regulates longitudinal sheath stability in oligodendrocytes.

Our work unexpectedly revealed that the process of sheath accumulation and stabilization involves extensive initial sheath loss. However, this contrasts with previous studies reporting that oligodendrocytes maintain most of the sheaths that are formed (5, 8). Those studies were the first to demonstrate that oligodendrocytes accumulate sheaths over the course of a few hours. However, imaging intervals of 10-20 minutes were used, and sheaths had to be at least 5 μm in length and stable for > 10 minutes to be counted. Our studies show that immature ensheathments as short as 2 μm can wrap around axons, and many of these disappear before growing to 5 μm. To detect these initial ensheathments, 5 minutes was the minimum imaging interval based on our criteria. Our work builds on these earlier studies (5, 8) to show that while oligodendrocytes accumulate sheaths in a relatively short time, this phase is highly dynamic and involves extensive sheath initiation and loss.

Myelin sheath formation can be most simply described as a combination of axonal adhesion and membrane biogenesis. So why are so many immature ensheathments destabilized and lost? Our axonal ensheathment dynamics experiment demonstrates that oligodendrocytes will ensheath the same domain of an axon an average of ~3 times before a stable sheath is formed. We propose that this extensive “sampling” of axons finds targets that are ready to be myelinated, or that this sampling may modify axons to enhance their ability to be myelinated. Successful sheath formation presumably occurs on axons that are primed with the appropriate adhesion molecules needed to facilitate binding and downstream wrapping. Previous work shows that the disruption of different adhesion molecules in both neurons and oligodendrocytes can result in reduced myelin sheath number (22–25). To further build on this concept, axonal vesicular release could be a part of this priming mechanism. This is suggested by the observation that synaptic proteins in axons accumulate underneath and around myelin sheaths to promote stability and sheath growth (7, 8, 24, 26). Interestingly, recent evidence suggests that myelin stimulates this vesicular release as part of a feed-forward mechanism to further promote myelination (26). It is tempting to speculate that repetitive ensheathment attempts on the same domain of an axon could be part of a mechanism for priming axons so that stable sheaths can be formed.

Our work supports a model where Rab5+, Rab7+, and Rab11+ endocytic endosomes localize to immature ensheathments and that Rab5 regulates the long-term stabilization of sheaths. It seems reasonable that the endocytic recycling pathway could be regulating the localization of adhesion proteins within myelin to help optimize sheath stability. The decrease in sheath stability that we observe when inhibiting Rab5 is consistent with some of the phenotypes reported from disrupting adhesion proteins in myelin (22–25). Previous work showing that the endocytic recycling pathway is critical for the proper localization of myelin proteins such as PLP in oligodendrocyte cultures also supports this idea (27–31). Our study could suggest that the recycling of adhesion molecules or other myelin proteins is most prevalent during the stabilization phase of sheath formation. However, since we over-expressed a dominant-negative mutant to inhibit Rab5 activity using *myrf*,which is a specific but weak driver, it is possible that the Rab5 expression levels were insufficient to disrupt the early accumulation phase of sheath formation. It seems reasonable that the recycling of adhesion proteins would also be important during this phase as oligodendrocytes ensheath the same axons multiple times before stabilizing any sheaths.

It is very intriguing that there are regional differences in the number of ensheathments formed by each oligodendrocyte, yet the stabilization rate is relatively consistent across all cells. Dorsal oligodendrocytes are less abundant but produce more ensheathments per cell than ventral oligodendrocytes. Despite these differences, cells from both regions exhibit a similar ratio of ensheathments formed to ensheathments lost. The low rate of stabilization was very similar in two different experimental paradigms, further substantiating this conclusion. We analyzed ensheathment dynamics on individually labeled axons (Figure 2) and across individual oligodendrocytes (Figures 3–5). These showed that ~21% and ~15% (respectively) of the total number of ensheathment attempts stabilized. This is clearly a fundamental parameter of the sheath formation process since it is so consistent across all cells.

Could this consistent stabilization rate be an intrinsic property of oligodendrocytes? Given the complexity of the zebrafish *in vivo* system, that is difficult to conclude. However, it is possible that oligodendrocytes balance total ensheathment attempts with the rate of stabilization based on the size of the axons that are being wrapped. Cells wrapping large caliber axons might not be able to support as many total sheaths due to the increased amount of membrane required to wrap larger axons. The mechanism by which oligodendrocytes would sense the caliber of each axon is still speculative. However, this study could suggest that oligodendrocytes repetitively ensheath the same axons as a means of detecting the diameter. Each wrapping attempt could involve a different amount of membrane, allowing the oligodendrocyte to use the physical sites of axonal contact during each attempt as a guide for subsequent myelination.

This study highlights several interesting observations regarding how oligodendrocyte sheath number is regulated at both the cellular and molecular level. There is a clear relationship between the number of ensheathment attempts performed by an oligodendrocyte and the number of these ensheathments that are ultimately stabilized during development. While the endocytic recycling pathway helps to promote the stability of sheaths, this process is still relatively unexplored. Therefore, this work will lead to important future research examining how the mechanisms regulating membrane biogenesis and sheath stability may cooperate to determine sheath number vs. sheath thickness and length.

## Materials and methods

All data and reagents will be made available upon reasonable request to Dr. Wendy Macklin.

### Zebrafish lines and husbandry

The Institutional Animal Care and Use Committee at the University of Colorado School of Medicine approved all animal work (#00419). This group follows the U.S. National Research Council’s Guide for the Care and Use of Laboratory Animals and the U.S. Public Health Service’s Policy on Humane Care and Use of Laboratory Animals. Zebrafish larvae were raised at 28.5°C in embryo medium and were staged as days post fertilization (dpf). Transgenic lines used in this study were *Tg(nkx2.2a:EGFP-CAAX), Tg(mbp:eGFP-CAAX*), and *Tg(mbp:tagRFP*) (32–35). All other constructs were introduced by transient transgenesis to achieve sparse labeling for single-cell analysis in wild type ABs. All experiments and analyses were performed blind to gender since sex is determined at later stages than what was studied here.

### Plasmid construction

Multi-site gateway cloning (36) was used to produce the following plasmids: *pEXPR-sox10:eGFP-CAAX, pEXPR-sox10:mScarlet-CAAX* (gift from Bruce Appel’s lab)*, pEXPR-neuroD:tagRFP-CAAX*, and *pEXPR-mbp:eGFP*. The first 3 plasmids contain a cysteine-aliphatic amino acid-X (CAAX) prenylation motif that targets the fluorescent protein to the cell membrane. The following entry plasmids were used to build these constructs: *p5E-sox10* (37), *p5E-neuroD* (gift from Bruce Appel’s lab), *p5E-mbp* (35), *pME-eGFP-CAAX, pME-tagRFP-CAAX*, *p3E-eGFP, p3E-polyA*, *pDEST-Tol2* (no transgenesis marker).

The *pEXPR-mbp:eGFP* plasmid was linearized using *SalI* and *SnaBI* to remove both the *mbp* driver and *eGFP* insert. An *myrf* driver element was PCR amplified from *p5E-myrf* (gift from Bruce Appel’s lab) and was ligated into the linearized vector with an *eGFP* insert flanked with *BamHI* and *Age*I sites using NEBuilder^®^ HiFi assembly cloning. The resulting *pEXPR-myrf:eGFP* plasmid was confirmed by diagnostic digest and sequencing and was used as a base vector for building the remaining over-expression constructs.

Plasmids encoding the *RAB5C, RAB7A*, and *RAB11A* zebrafish coding sequences (Addgene, *RAB5C* = 80518, *RAB7A* = 80522, and *RAB11A* = 80529, (38)) were sequenced, and the *RAB7A* and *RAB11A*sequences each had a point mutation *(D63G* and *Q166R* respectively) relative to the NCBI protein sequences *(RAB7A* =NM_200928.1, *RAB11A* =NM_001007359.1) and relative to other previous work in zebrafish (19). These mutations were corrected using a combination of mutagenesis by PCR-driven overlap extension (39) and NEBuilder^®^ HiFi assembly cloning.

The *RAB5C, RAB7A*, and *RAB11A* sequences were then cloned directly into the *pEXPR-myrf:eGFP*plasmid as follows. This plasmid was linearized using *BamHI* and *AgeI* to remove the *eGFP* insert. Each *RAB* sequence was PCR amplified using the following primer sequences (additional overhang sequences were added to each primer for NEBuilder® HiFi assembly-based cloning that are not shown here):

- atggcggggcgaggtgg (Rab5c F)
- ttagtttccgcctccacagc (Rab5c R)
- atgacatcaaggaagaaagttcttctgaagg (Rab7a F)
- tcagcagctacaggtctctgc (Rab7a R)
- atggggacacgagacgacg (Rab11a F)
- ctagatgctctggcagcactgc (Rab11a R)

The *RAB* PCR fragments were then ligated into the linearized plasmid, along with either *tagRFP* or *eGFP* PCR fragments, using NEBuilder® HiFi to generate the following fusion constructs (a kozak sequence, GCCACC, was added directly 5’ of the translational start site for all constructs):

- *pEXPR-myrf:eGFP-RAB5C*
- *pEXPR-myrf:eGFP-RAB7A*
- *pEXPR-myrf:eGFP-RAB11A*
- *pEXPR-myrf:tagRFP-RAB5C*
- *pEXPR-myrf:tagRFP-RAB7A*
- *pEXPR-myrf:tagRFP-RAB11A*
- *pEXPR-myrf:tagRFP*

We also made a set of dominant-negative point mutations for each of the *RAB* genes *(rab5C^S36N^, rab7A^T22N^, rab11A^S25N^*) that have been characterized in zebrafish previously (19). We generated the *rab5C^S36N^* mutant through a combination of mutagenesis by PCR-driven overlap extension (39) and NEBuilder® HiFi assembly cloning. Primer sequences:

- *rab5C^S36N^*
  - PCR fragment 1, mutation site at 3’ end of R2 primer.
    - atggcggggcgaggtgg (F)
    - cagactctcccagcaacacaagc (R1)
    - ccaggctgttcttgcctactgcagactctcccagcaacacaagc (R2)
  - PCR fragment 2, mutation site at 5’ end of F2 primer.
    - tgctgcgcttcgtcaaaggc F1
    - cagtaggcaagaacagcctggtgctgcgcttcgtcaaaggc (F2)
    - ttagtttccgcctccacagc (R)

Both final PCR fragments were then ligated into the same linearized *pEXPR:myrf* plasmid described above, along with a *tagRFP* PCR fragment, using NEBuilder® HiFi to generate the following fusion plasmid (a kozak sequence, GCCACC, is directly 5’ of the translational start site):

- *pEXPR-myrf:tagRFP-rab5C^S36N^*

The *rab7A^T22N^* and *rab11A^S25N^* mutations were made by Keyclone Technologies (San Marcos, CA). Resulting plasmids:

- *pEXPR-myrf:tagRFP-rab7A^T22N^*
- *pEXPR-myrf:tagRFP-rab11A^S25N^*

All plasmids were confirmed by diagnostic restriction digest and sequencing.

### Injections and general imaging parameters

Plasmids were injected into zebrafish embryos at the single-cell stage with Tol2 mRNA to achieve transient transgenesis and sparse labeling. On the desired day, the injected larvae were mounted laterally in 0.8% low-melt agarose with 140 ug/mL (0.014%) of Tricaine as anesthesia. All imaging was performed live using a Nikon A1R resonance scanning confocal microscope and a 40X Apochromat long-working distance water immersion objective with a 1.15 NA (Nikon MRD77410). A temperature-controlled stage maintained at 28.5°C was used for all time-lapse experiments. All imaging was done in the spinal cord of each larva above the yolk-sac extension. Only a single cell was imaged from each larva unless otherwise noted in each figure legend. Oligodendrocytes in the ventral spinal cord that myelinated Mauthner axons were excluded from all analyses in this work.

### Myelinating oligodendrocyte cell counts in the spinal cord

*Tg(mbp:eGFP-CAAX*) and *Tg(mbp:tagRFP*) transgenic lines that express membrane tethered eGFP and cytosolic tagRFP in myelinating oligodendrocytes respectively were crossed and embryos were grown to 4dpf. Both sides of the spinal cord for each larva were imaged laterally above the yolk-sac extension with a 1X optical zoom (0.31 μm XY pixel size), a 0.3 μm z-step size (Nyquist), and 32X line averaging. Cell counts were performed in Imaris (version 9.8). Each image was cropped to separate dorsal and ventral regions. A gaussian filter was then applied with a 0.311 μm width to smooth out noise. Following this, a local-background subtraction was performed with an estimated cell body size of 7 μm. A threshold for fluorescent intensity was then applied to each image manually. Finally, a 0.5 μm water-shed filter was used to separate contacting cell bodies and Imaris counted the number of cells in each image. Cell bodies occupying the area in between the dorsal and ventral axon tracts were manually removed (these cells were in the minority).

### Static oligodendrocyte imaging for sheath number/length analysis

Individual sheaths were visualized by labeling oligodendrocytes with *sox10:eGFP-CAAX* by transient transgenesis. These cells were imaged at 4 dpf with a 2X optical zoom (0.16 μm XY pixel size), a 0.3 μm z-step size (Nyquist), and 32X line averaging. Cells were chosen that were not too crowded for accurate quantification. Except for Figure 1D, dorsal and ventral cells were never combined for this analysis. In each image every sheath was identified and counted by inspecting individual optical sections. The lengths of each sheath were measured from maximum intensity projections using the simple neurite tracer plugin in Fiji. From this data we calculated sheath number, average sheath length, and the total sheath length per cell (calculated by adding up the length of all sheaths supported by each cell).

### Axonal ensheathment dynamics imaging and analysis

Criteria for quantifying ensheathment attempts using labeled axons were established. Axon membranes were sparsely labeled with the *neuroD:tagRFP-CAAX* plasmid by transient transgenesis in the *Tg(nkx2.2:eGFP-CAAX*) stable line. At 2.5dpf larvae were anesthetized in tricaine and mounted, and time-lapse imaging was performed for 15-18 hours with an imaging interval of 5 minutes. A 2X optical zoom (0.16 μm XY pixel size), a 0.5 μm z-step size, and 16X line averaging were used. Imaging was done in the spinal cord above the yolk sac extension and focused on dorsal and midline axons, since the *Tg(nkx2.2:eGFP-CAAX*)line was too crowded in the ventral region for visualizing ensheathment dynamics. Regions in each video with potential axonal ensheathments were cropped and corrected for drift using the 3D drift correct plugin in Fiji. Individual optical sections were inspected and a volume projection in Imaris was analyzed for each axonal ensheathment. From this, we developed quantification criteria. Ensheathments were ~2 μm or longer and cylindrical in shape around the axon, i.e., the axon fluorescent signal had to go through the center of the oligodendrocyte signal, and it had to be more than ~75% of the way around the axon. Once an ensheathment had formed, it had to shrink to below 2 μm in length and be less than half-way around the axon to be considered lost. Ensheathments were considered destabilized if they were lost by the end of the time-lapse period and stable if they remained.

Repetitive ensheathments were multiple rounds of sheath initiation/loss on the same domain of an axon. Each axon domain was the region between the two most lateral ensheathment attempts that took place during a series of repetitive ensheathments. In cases with only a single ensheathment attempt, the axon domain was the region directly underneath the sheath. These domains ranged from ~2-14 μm in length. Some axons had more than one domain that was ensheathed. These domains were considered separate because they were ensheathed by different oligodendrocytes or because a single oligodendrocyte had more than one sheath on the same axon at the same time. Out of 25 axonal domains, 18 were repetitively ensheathed more than once and each of these ensheathment attempts were always from the same oligodendrocyte. Fourteen of these domains were repetitively ensheathed by the same process extending from each oligodendrocyte.

### Oligodendrocyte ensheathment dynamics imaging and analysis

Oligodendrocytes were sparsely double-labeled with our *pEXPR-sox10:eGFP-CAAX* plasmid and one of the following plasmids using transient transgenesis: *pEXPR-myrf:tagRFP, pEXPR-myrf:tagRFP-RAB5C*, or *pEXPR-myrf:tagRFP-rab5C^S36N^*. The *myrf* driven plasmids identified cells that were fated to make myelin. At 2.5dpf, larvae were anesthetized in tricaine and mounted, and eGFP+/RFP+ progenitor cells were chosen for imaging above the yolk-sac extension in the spinal cord. Time-lapse imaging was performed for 15 hours with an imaging interval of 5 minutes. A 2X optical zoom (0.16 μm XY pixel size), a 0.75 μm z-step size, and 16X line averaging were used. After the time-lapse imaging period was over at 3 dpf, the larvae were removed from the mounting agar and were put back into embryo medium in separate wells of a 24-well plate. At 4 dpf, a final static image was taken for each of the same cells to conclude the imaging paradigm. To be considered for analysis, we had to obtain at least 30 minutes of imaging before each cell started the ensheathment process. Additionally, each cell had to stop making sheaths for at least 90 minutes before the end of the imaging interval.

A max projection of each video was background subtracted and corrected for drift using Fiji’s 3D drift correct plugin. Sheath initiation and loss were counted manually in every frame using the multi-point counter in ImageJ based on the ensheathment criteria we established in the previous section. Although the counter ROIs for this were placed within each max projection, every sheath initiation/loss event was determined by inspecting individual optical sections and by looking at a volume projection in Imaris for each frame. At the final 4 dpf time point, all sheaths were counted, and the lengths were measured using the simple neurite tracer plugin in Fiji.

A quality control experiment determined that the conditions of this imaging paradigm did not significantly change the average sheath number or sheath length across a population of cells. Static images of both dorsal and ventral oligodendrocytes were collected at 4 dpf from zebrafish larvae that had been anesthetized in tricaine, mounted in agar, and housed in the time-lapse imaging chamber for the 15-hour experiment from 2.5-3 dpf. However, these larvae were not imaged during this time. We compared the sheath number and lengths for these cells (agar_tricaine condition) to our sheath number counts from Figure 1c (standard condition) and to cells that went through the full ensheathment dynamics imaging paradigm (time-lapse condition) (Supp figure 3.1a-d).

A quality control experiment determined that oligodendrocytes did not make any new ensheathments during the stabilization phase from 3-4dpf. Cells were imaged using a modified ensheathment dynamics imaging paradigm. First, from 2.5-3dpf, the imaging interval was 30 minutes rather than 5 minutes. At 3dpf, the larvae were removed from the tricaine and imaging agar and were allowed to recover in embryo media for 30 minutes. After this the larvae were remounted, and the same cells were imaged for an additional 23 hours with an imaging interval of 15 minutes. We tracked sheath initiation and loss from 3-4dpf for 5 dorsal cells and 3 ventral cells. None of these 8 oligodendrocytes produced any new sheaths from 3-4dpf (Supp fig 5.1).

### Endosome imaging and quantification

Our *myrf:eGFP-RAB5C, -RAB11A*, and *-RAB7A* fusion constructs were transiently expressed alongside *sox10*:mScarlet-CAAX to label the membrane of individual oligodendrocytes. At 2.5dpf, larvae were anesthetized in tricaine, mounted, and cells that were in the early stages of the ensheathment process were chosen for imaging. Static snapshots of a mix of both dorsal and ventral cells were taken using a 3X optical zoom (0.1 μm XY pixel size, Nyquist), a 0.75 μm z-step size, and 16-32X line averaging. A larger z-step size than Nyquist (0.3 μm) was used to minimize motion artifacts during imaging. Images were pre-screened by viewing the membrane marker channel only and individual sheaths with high-quality imaging were cropped for further processing. The number of endosomal puncta in each image was quantified using a spots analysis in Imaris (version 9.2). We used an estimated XY diameter of 0.5 μm to detect Rab5 and 0.4 μm to detect Rab7 and Rab11. After a local-background subtraction, a threshold for fluorescent intensity was applied to each image manually. Imaris counted the number of puncta, and we normalized these numbers to the length of each sheath (in microns), measured using the Imaris filament tracer. Only puncta within a sheath were counted based on overlap with the membrane marker. Puncta that were present in adjacent processes or other structures were manually excluded from each image.

### Statistics, sample size determination, and reproducibility/rigor

All statistics and plots were done using Graphpad Prism (version 9). We used the non-parametric Mann-Whitney test for all unpaired comparisons with no assumption of normality. For groups of 3 or more, global significance was first assessed using the non-parametric Kruskal-Wallis test. This was followed up by Dunn’s multiple comparison tests if the global P value was <0.05 (unless otherwise stated in each figure legend). Global P-values are reported when not significant. Non-parametric statistical tests were chosen since each data set had groups that were not normally distributed and/or had unequal variances. Individual data points are shown for all plots with the central dashed lines representing the average of the group. We considered P < 0.05 the threshold for statistical significance and we provide exact P values for all analyses.

The sample sizes for the cell counts in Figure 1, the repetitive axonal ensheathment analysis in Figure 2, the Rab+ endosomal quantification in Figure 6, and the modified ensheathment dynamics data in supplemental figure 5.1 were not pre-determined. The sample sizes for the sheath analyses performed in Figures 1, 7 and supplemental figure 3.1 were determined based on previous practices and are meant to show the full distribution for this type of data set. The control, *RAB5*, and *rab5C^S36N^* ‘sheath number per cell’ data in Figure 7 was used to do a power analysis (GPower Version 3.1) to determine the appropriate sample sizes for the ensheathment dynamics data in Figures 3–5, 8, and supplemental figures 3.1 (time-lapse group) and 8.1. We performed a power analysis using the parametric “ANOVA: Fixed effects, omnibus, one-way” stats test in GPower with 3 groups (control, *RAB5, rab5C^S36N^*) with an alpha error probability of 0.05 and 80% power. The effect size was calculated within GPower. From this, GPower calculated needing 15 cells per group. Since sheath number was not normally distributed, we had to assess statistical significance for these experiments using a non-parametric test. We therefore increased the sample size by 20% (n = 15 increased to n = 18) to account for the decreased power of non-parametric tests. The same time-lapse ventral control data set was used in Figures 4, 5, 8, and supplemental Figure 3.1 because this data is very difficult to collect, and the analysis was very time intensive. Therefore, we also used the results of this power analysis to determine the same sample size of the dorsal group of cells analyzed in Figures 3–5 and supplemental Figure 3.1 (time-lapse group).

All representative images in each figure were from the associated quantified data set. These images were adjusted for brightness and contrast for clarity of presentation. No quantifiable data was excluded from any analysis. All data in each figure was pooled from a minimum of 2 independent rounds of fish crossing and injections. Data for each individual cell in this study is considered a biological replicate. Separate data points from the same cell (for example: different sheath lengths from the same cell) are considered as technical replicates in this study. All analyses were performed blind using a custom Fiji script called “blind-files” (source code can be found at https://github.com/Macklin-Lab/imagej-microscopy-scripts).

## Acknowledgements

This work was supported by National Institutes of Health R37NS82203 to W.B.M. and NIH 5F31NS118830 to A.R.A. We thank Dr. Andrew Lapato and Dr. Caleb Doll for valuable discussion and feedback on this manuscript.

## Author contributions

A.R.A and W.B.M. designed the research.

A.R.A. performed the research, analyzed the data, and wrote the manuscript

A.R.A. and W.B.M. edited the manuscript.

---

**Fig. 3.1 supp.**
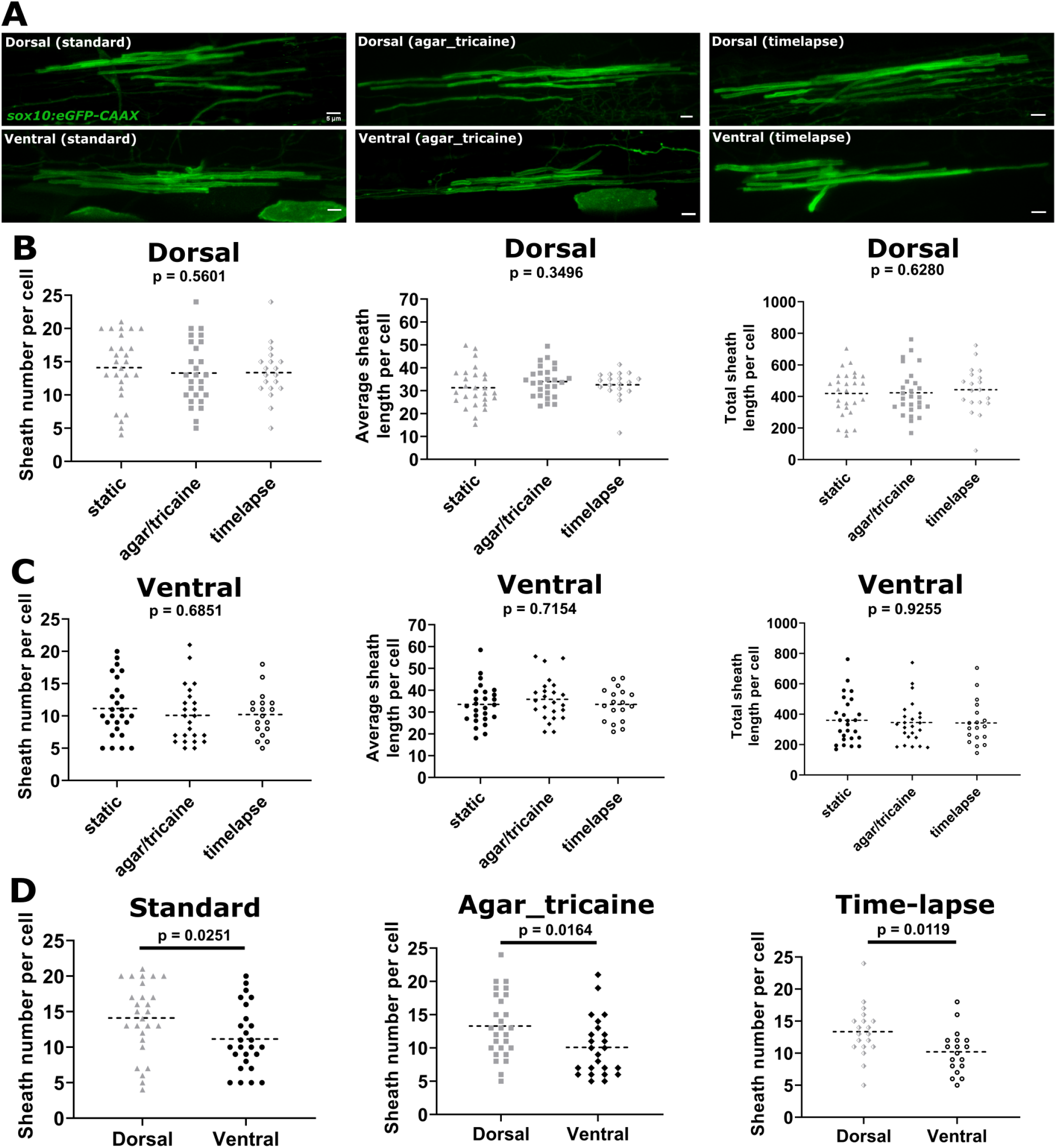
Ensheathment dynamics imaging conditions did not significantly alter sheath number or length. **A**) Representative lateral images of dorsal and ventral oligodendrocytes for each of three testing groups (standard, agar_tricaine, time-lapse) in the spinal cord of living larvae at 4dpf labeled by *sox10:eGFP-CAAX* (myelin in green). The images and data for the standard group are the same as in Figure 1. (Scale bar = 5 μm). **B**) Sheath number per cell, average sheath length per cell, total sheath length per cell from left to right for all dorsal data sets. **C**) Sheath number per cell, average sheath length per cell, total sheath length per cell from left to right for all ventral data sets. **D**) Sheath number per cell comparing the dorsal and ventral populations within each testing group. Standard: dorsal n=27 cells/27 larvae, ventral n=26 cells/26 larvae; agar_tricaine: dorsal n=24 cells/24 larvae, ventral n=25 cells/25 larvae; time-lapse: dorsal n=19 cells/19 larvae, ventral n=18 cells/18 larvae. The dashed lines in each plot represent average values with all data points shown. Significance determined by Kruskal-Wallis test for (B, C). Global p-values are presented for each of these plots. Post hoc multiple comparisons tests were not performed since the global p-values were not significant. Mann–Whitney tests were performed for (D).

**Fig 3.2 supp.**
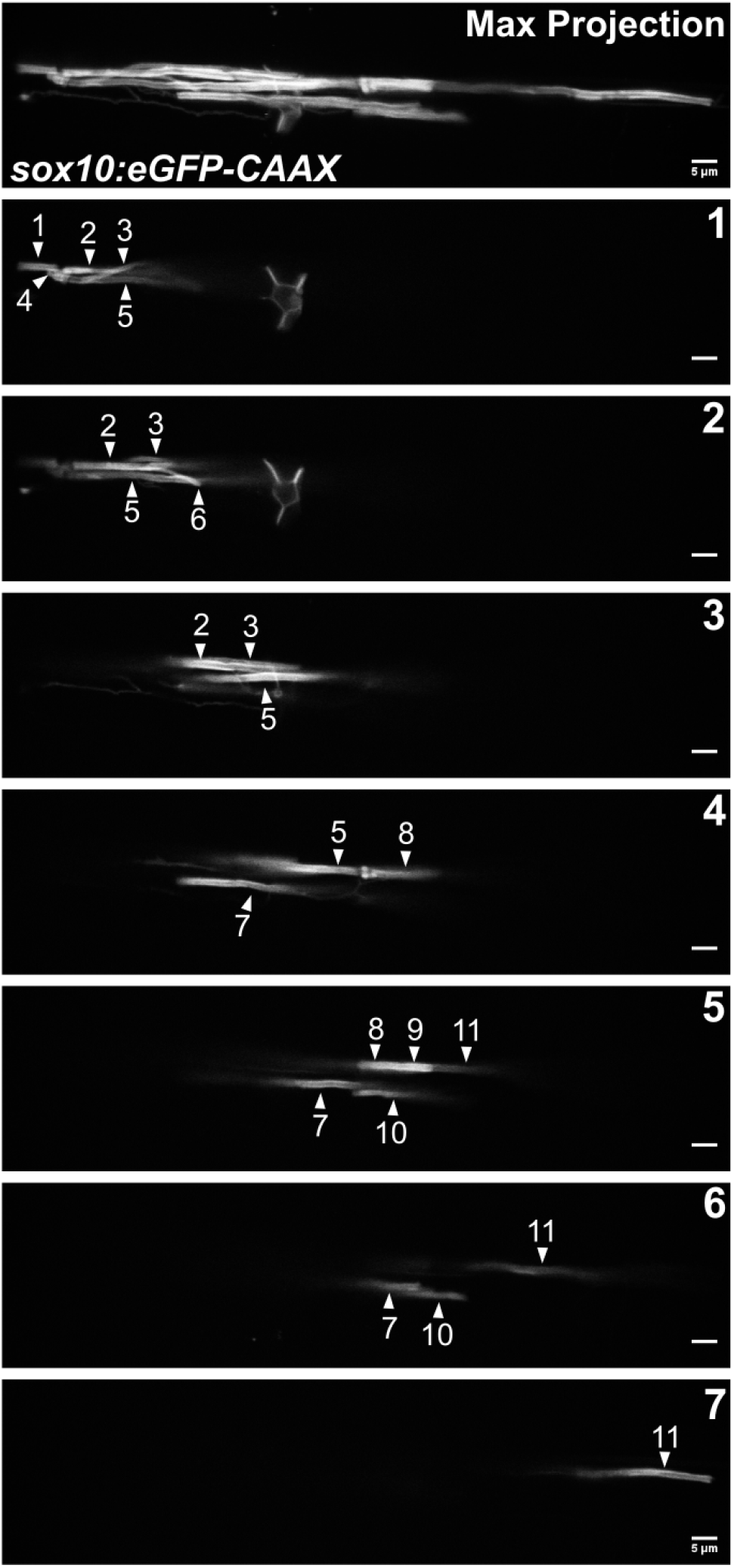
Visual quantification of sheath number at 4dpf for the representative cell in Figure 3C. Lateral images of an oligodendrocyte in the spinal cord of living larvae labeled with *sox10:eGFP-CAAX* at 4dpf. The first panel is a max projection image of the entire cell. Each subsequent panel is a single optical section labeled 1-7. White arrows are pointing out 11 different sheaths. (Scale bar = 5 μm).

**Fig 5.1 supp.**
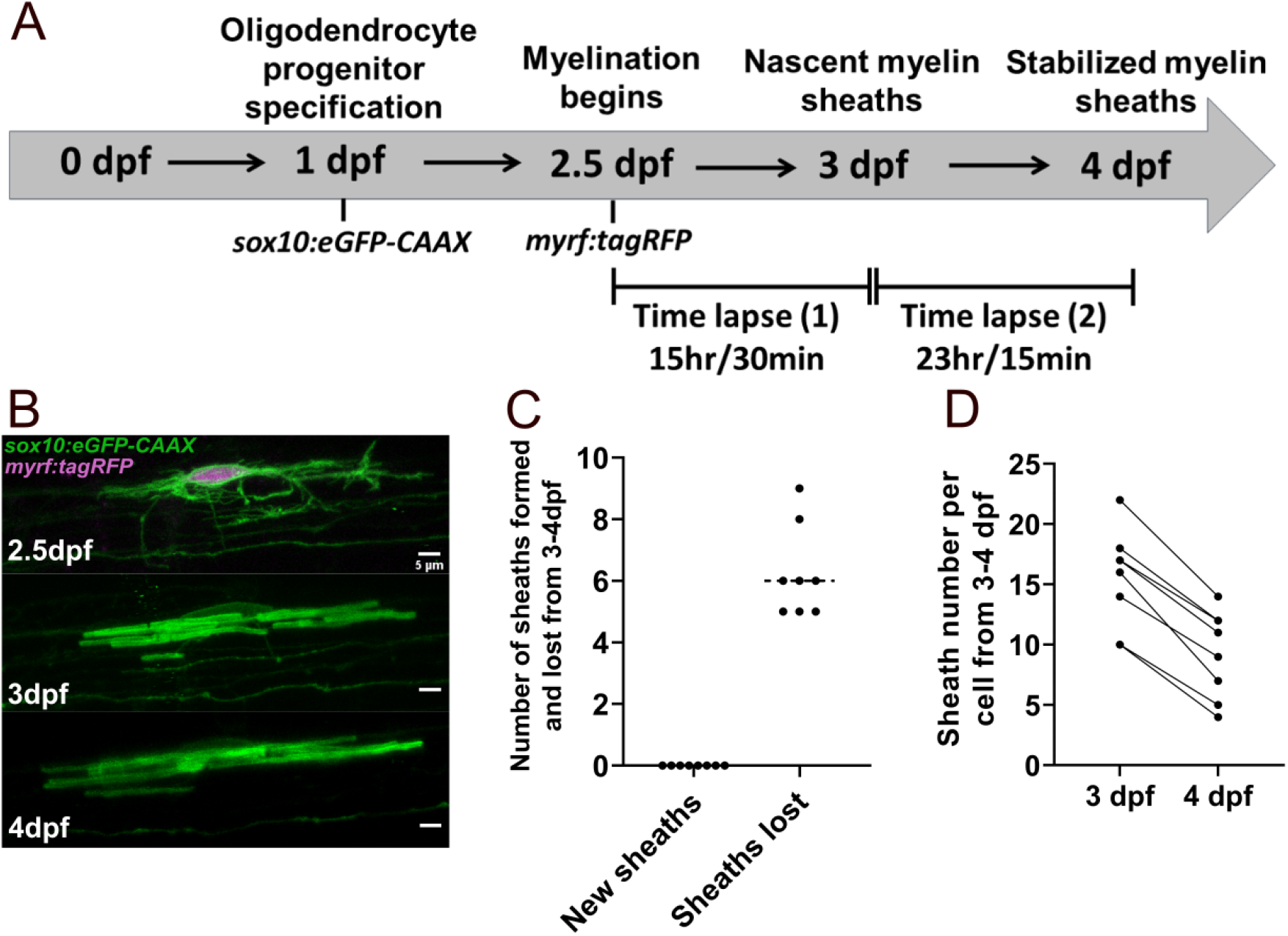
Oligodendrocytes do not make new sheaths during the stabilization phase from 3-4dpf. **A**) Modified oligodendrocyte ensheathment dynamics imaging paradigm. **B**) Lateral images in the spinal cord of living larvae labeled with *sox10:eGFP-CAAX* (in green) and time-lapsed from 2.5-4dpf. The upper panel is an oligodendrocyte that is also labeled with *myrf:tagRFP* (magenta) at the beginning of the time-lapse experiment. The subsequent panels are of the same cell at 3dpf and 4dpf. (Scale bar = 5 μm). **C**) Number of sheaths formed and lost from 3-4dpf per cell. The dashed lines in this plot represent average values with all data points shown. **D**) Change in sheath number from 3-4dpf per cell. The straight lines are connecting the number of sheaths at 3dpf and 4dpf for each cell. Dorsal n=5 cells/4 larvae, ventral n=3 cells/3 larvae.

**Fig. 8.1 supp.**
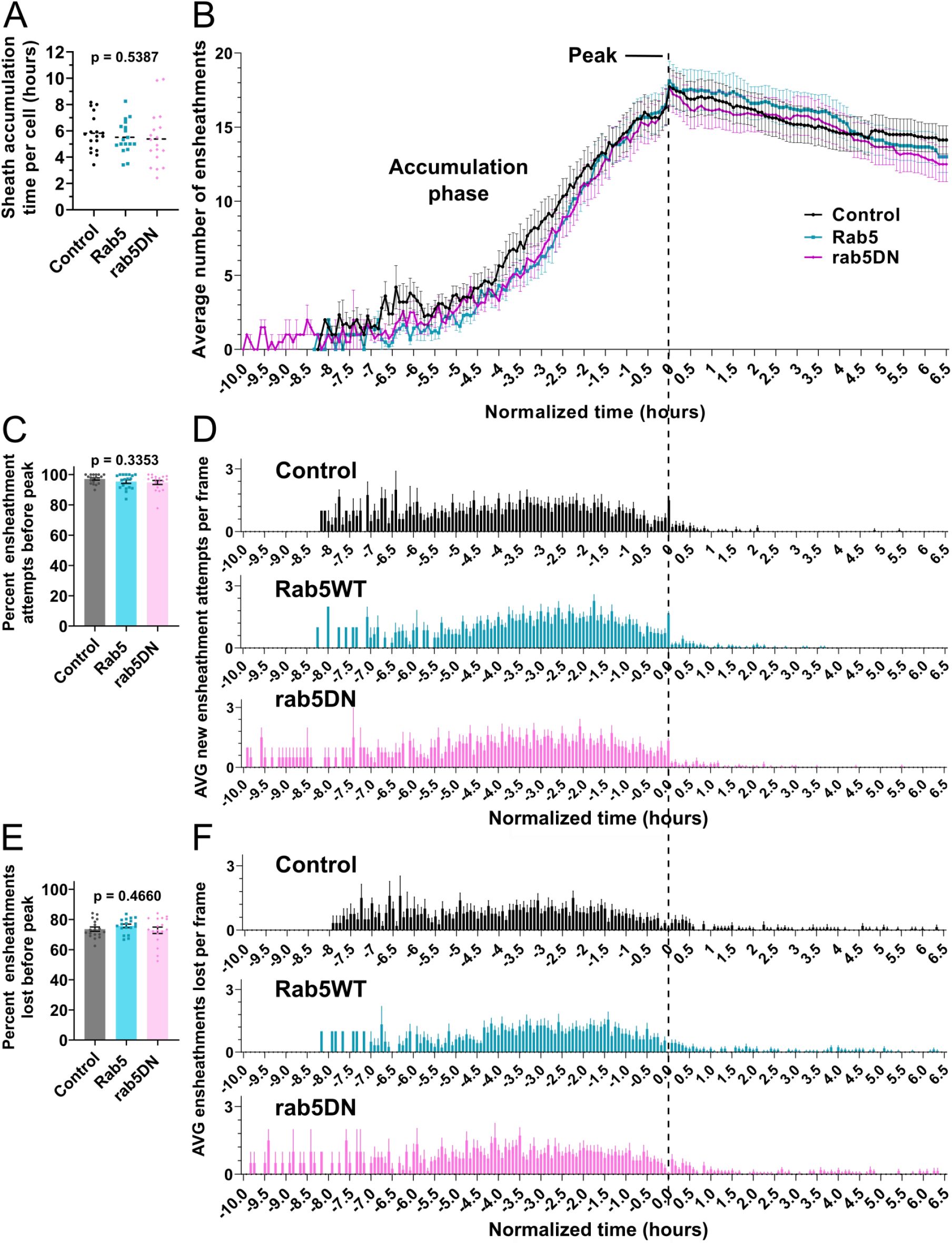
The dynamics of the sheath accumulation phase were not altered by *rab5C^S36N^* over-expression. **A**)Time required for peak (or max) sheath accumulation per cell (hours). **B-F**)The control, *RAB5C*, and *rab5C^S36N^* time-lapse data sets were quantified for sheath initiation and loss in each 5 minute frame. This data was normalized by setting the video frame where each cell accumulates a peak number of immature sheaths as T0’, as in Figure 4. The vertical dashed line aligns the T0’ time point of each graph in B, D, and F. **B**) The average number of immature ensheathments in each frame are plotted as a line with error bars representing SEM. The data for the control is the same as for the ventral group in Figure 4. **C**) Percent of the total ensheathment attempts that occurred prior to reaching peak. Error bars represent SEM. **D**) The average number of new ensheathment attempts in each frame was quantified with the same time normalization as in B. Error bars represent SEM. **E**) Similar as in C, the percent of the total number of immature ensheathments that were lost prior to reaching peak. Error bars represent SEM. **F**) The average number of immature ensheathments that were lost in each frame was quantified with the same time normalization as in B. Error bars represent SEM. Significance was determined using global Kruskal-Wallis tests in A, C, and E. These p-values are shown for each plot since they were not significant. Post hoc multiple comparisons tests were not performed for these analyses. Control n=18 cells/18 larvae, wild-type *RAB5C* n=18 cells/18 larvae, *rab5C^S36N^* n=18 cells/17 larvae. The data in B, D, and F was cropped at +6.5 hours.

